# Dynamics of DNA replication during male gametogenesis in the malaria parasite *Plasmodium falciparum*

**DOI:** 10.1101/2021.12.18.473304

**Authors:** Holly Matthews, Jennifer McDonald, Francis Isidore G. Totanes, Catherine J. Merrick

## Abstract

Malaria parasites undergo a single phase of sexual reproduction in their complex lifecycle, during which they cycle between mosquito and vertebrate hosts. Sexual reproduction occurs only at the point when parasites move into the mosquito host. It involves specialised, sexually committed cells called gametocytes, which develop very rapidly into mature gametes and then mate inside the mosquito midgut. The gamete development process is unique, involving unprecedentedly fast replication and cell division to produce male gametes. A single male gametocyte replicates its ∼23Mb genome three times over to produce 8 genomes, segregates these into newly-assembled flagellated gamete cells and releases them to seek out female gametes, all within ∼15 minutes. Here, for the first time, we use fluorescent labelling of *de novo* DNA synthesis to follow this process at the whole-cell and single-molecule levels, yielding several novel observations. Firstly, we confirm that no DNA replication occurs before gametogenesis is triggered, although the origin recognition complex protein Orc1 is abundant even in immature gametocytes. Secondly, between repeated rounds of DNA replication there is no detectable karyokinesis – in contrast to the repeated replicative rounds that occur in asexual schizonts. Thirdly, cytokinesis is clearly uncoupled from DNA replication, and can occur even if replication fails, implying a lack of cell cycle checkpoints. Finally the single-molecule dynamics of DNA replication are entirely different from those in asexual schizonts.

## INTRODUCTION

The malaria parasite *Plasmodium* is one of the most successful human pathogens, infecting hundreds of millions of people every year [1]. It pursues a complex lifecycle involving massive asexual replication in both human and mosquito hosts, plus one phase of sexual reproduction in the mosquito. This is a key bottleneck in the *Plasmodium* lifecycle and an attractive target for transmission-blocking drugs or vaccines [2].

For sexual reproduction, immature gametes called gametocytes develop from intraerythrocytic *Plasmodium* parasites circulating in the human bloodstream. Only when they switch hosts into the gut of a blood-feeding mosquito do these gametocytes mature into male (micro) or female (macro) gametes and mate via fertilisation. The molecular and cellular biology of this process is highly unusual. Sexual cells do not undergo conventional binary fission [3]: instead the male gametocytes differentiate and divide very rapidly upon entering the mosquito, undergoing dramatic changes in morphology and an 8-fold increase in ploidy before dividing to produce 8 mature flagellated gametes. This all occurs within less than 15 minutes, requiring each gametocyte to replicate its single ∼23Mb haploid genome 3-fold, followed by karyokinesis and cytokinesis [4, 5]. If the resultant male gametes encounter mature female cells (which have also exited their host erythrocytes and undergone morphological changes but not replication), then two gametes can fuse to form a diploid zygote. The zygote leaves the midgut to encyst on the midgut wall, proceeds through meiosis and ultimately regenerates a haploid genome in the oocyst. Asexual reproduction then recommences, producing as many as thousands of human-infective sporozoites per oocyst [6] (Fig 1).

**Figure 1:**
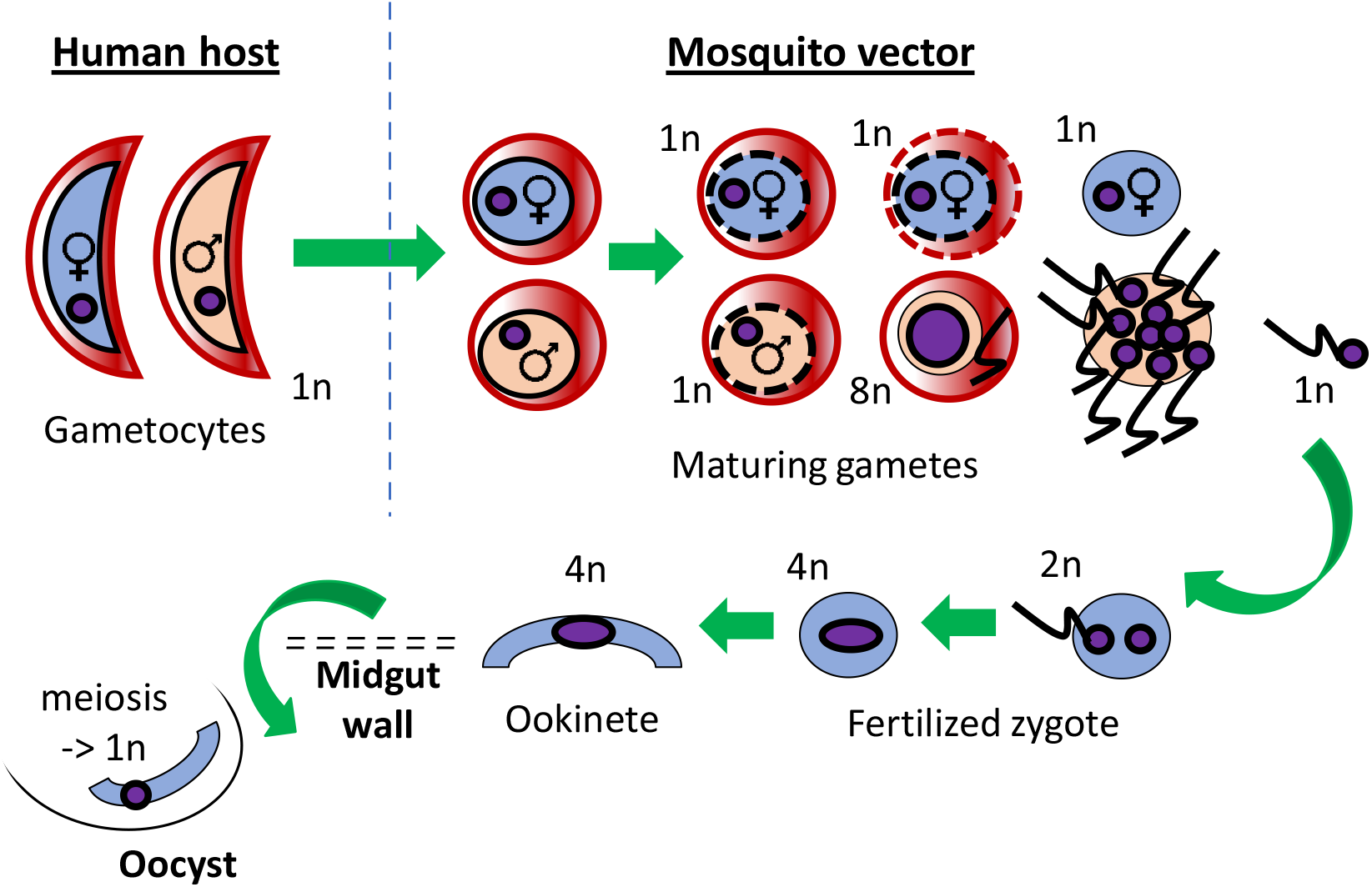
Schematic showing sexual development stages in *P. falciparum*. The progression of gametocytes to gametes, zygotes, ookinetes and oocysts is shown, highlighting phases of genomic replication that occur at each stage, giving cells between 1n and 8n genome contents.

The sequence of events shown in Figure 1 was described in seminal work conducted with rodent and bird malaria models [4, 5, 7] and the same processes occur with similar timing in human parasites such as *P. falciparum* [8, 9]. *P. falciparum* is, however, unusual compared to most other *Plasmodium* species in requiring a lengthy 2-week period to produce mature, crescent-shaped gametocytes in the human bloodstream.

The cellular phenomena are relatively well-studied but the underlying molecular biology is less so: the dynamics of gametocyte genome replication and the proteins controlling it remain incompletely understood. Male gametogenesis is clearly remarkable for its speed, demanding multiple genome replications at a pace unparalleled in eukaryotic gametogony – or indeed in any cellular replication process documented to date. Equally, the process is not analogous to any other model for non-binary-fission cell cycles, such as the syncytial replication seen in early *Drosophila* embryos. Here, many separate nuclei within a single cytoplasm undergo repeated binary divisions driven by synchronous waves of cyclin/CDK activity [10]. In *Plasmodium* gametogenesis, there is little evidence for conventional cycles of cyclin/CDK activity; indeed none of the cyclin-like proteins identified in *Plasmodium* appears to ‘cycle’ [11]. Furthermore, it is not clear whether replications and nuclear divisions are synchronous, or whether karyokinesis occurs at all prior to the final resolution into individual gametes, nor is it clear whether eight progeny must always be produced. Indeed, it is unclear why the parasite has evolved to replicate its male gametes at all. However, since gametocytogenesis is sex-biased with only 10-20% of cells being male (itself an unexplained bias [12]), the advantage of producing extra male cells to seek out scant female cells may outweigh the potential disadvantage of accelerated and error-prone replication.

Genome replication in *Plasmodium* has been best studied in erythrocytic schizogony – another atypical syncytial process. Schizogony requires hours not minutes per genome, with each replication and nuclear division in a schizont taking a total of 2-3 hours [13, 14]. Single-molecule measurements of DNA replication during schizogony revealed a mean replication-origin spacing of 65kb and a fork velocity of 1.2 kb/min [15], which is quite comparable to other eukaryotes. (Most human cell lines replicate at 0.5-2kb/min, while *S. cerevisiae* can achieve ∼2.9kb/min [16, 17]). To achieve gametogenesis, however, with genome replications completed in less than 5 minutes, a similar origin spacing would demand a fork velocity of at least 6.5kb/min – unprecedented in eukaryotes – or alternatively origins must be spaced at least every 12kb instead of 65kb. (A remarkably similar theoretical calculation was made in early work on *P. berghei* gametogenesis [18].) The latter is more likely, since extremely flexible origin usage has been documented in other organisms. In early *Xenopus* embryos, for example, replication occurs from origins spaced every 5-15kb, with origin density reducing after the mid-blastula transition [19, 20]. *Plasmodium* replication may be under equally flexible control, raising interesting questions about how flexible origin placement could be enforced in different lifecycle stages.

Here we have examined in detail the process of DNA replication in *P. falciparum* male gametogenesis, by exploiting a novel parasite line that incorporates a detectable modified nucleotide, bromodeoxyuridine (BrdU), into newly synthesised DNA [21]. We report that there is no replication before gametogenesis is triggered, although the origin recognition complex (an essential basal component of eukaryotic replication origins) appears to be present at high levels even in immature gametocytes. After triggering, there is no detectable karyokinesis during repeated rounds of replication; cytokinesis is uncoupled from DNA replication, implying a lack of cell cycle checkpoints; and finally the single-molecule dynamics of DNA replication are entirely different from those in asexual schizonts.

## RESULTS

### DNA replication in male gametocytes does not begin before exflagellation is activated

To measure *de novo* DNA synthesis during male gametogenesis, we first generated a new cell line in the NF54 strain of *P. falciparum*. The NF54 strain converts readily to gametocytes in culture, unlike the isogenic clonal reference strain, 3D7, which was previously used for our experiments on erythrocytic schizogony [15, 21]. NF54 parasites were transfected with a plasmid encoding thymidine kinase, which enables the parasites to salvage pyrimidine nucleosides, including thymidine analogs like BrdU [21]. We confirmed that this genetic modification did not affect differentiation into gametocytes and then proceeded to use this line for all future experiments.

Several reports in early literature suggested that male gametocytes begin the exflagellation process with DNA contents of ∼1.5n, markedly above the normal haploid content of 1n [4, 5, 8]. This would require some replication in the arrested gametocyte, but it has never been clarified at which stage in the 14-day maturation process this might occur. We exposed cultures of maturing gametocytes to a long (6h) pulse of BrdU at 8, 10, 12 and 14 days, i.e. stages III to V of maturation. Any cells synthesising DNA at these times should become labelled with BrdU, detectable at the cellular level by immunofluorescence (Fig 2A) and at the population level by ELISA (Fig 2B,C) [21]. DNA synthesis was never observed by immunofluorescence in a gametocyte at any stage examined in this study: the only cells incorporating BrdU at a population level were clearly rare asexual cells that survived in the gametocyte culture (Fig 2A) and these died off over time as expected, until they comprised only ∼1% of the total culture by day 14 (Fig 2C). Consistent with this, population-level BrdU incorporation remained extremely low throughout the timecourse, whereas a control asexual culture incorporated large amounts of BrdU (Fig 2B). Control ELISAs conducted in parallel showed that the developing gametocytes had synthesised large amounts of α-tubulin, as expected, whereas asexual cells contained relatively little.

**Figure 2:**
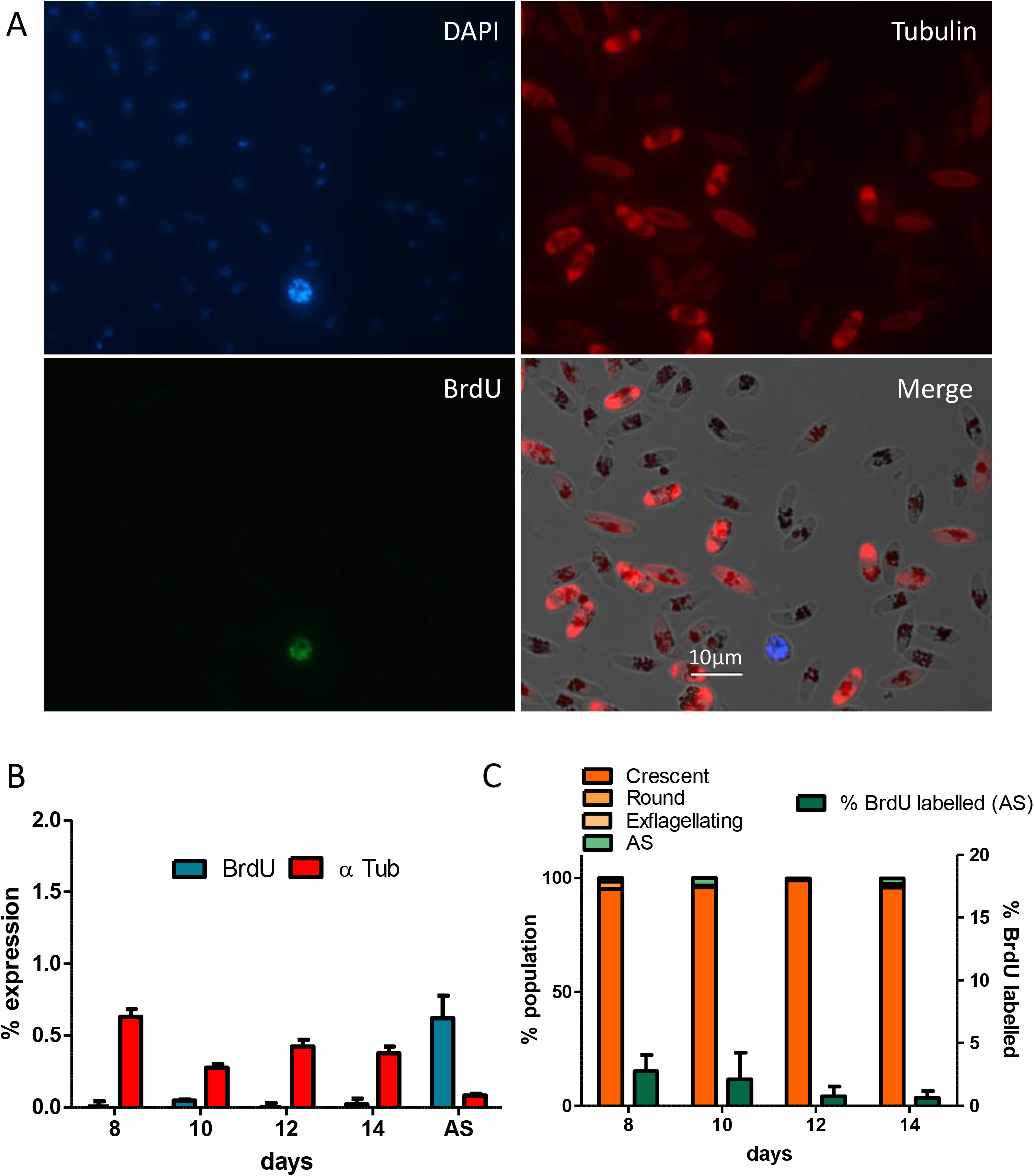
DNA synthesis measurements in unactivated gametocytes. Immature gametocyte cultures (pre-activation) were incubated with BrdU for 6 h and then analysed for incorporation of BrdU on days 8, 10, 12 and 14. **A)** Immunofluorescence images showing a large field of gametocytes at day 10, with no BrdU labelling, and a single asexual schizont incorporating BrdU. DAPI was included for alignment with BrdU labelling and for confirmation of morphology. **B)** Population-level ELISA measurement of DNA synthesis (BrdU) and α-tubulin levels (α-tub). All ELISA data represents the mean of at least three independent replicates, conducted in technical triplicate. An asexual culture was included as a control, showing high levels of BrdU incorporation and relatively low α-tubulin content. **C)** Single-cell-level immunofluorescence images (100 cells per timepoint) were categorised for cellular morphology (left axis) and for BrdU labelling (right axis). Data represents three independent counts of 100 parasites per timepoint.

Despite the absence of active replication prior to triggering, we found that the origin recognition complex subunit 1 (Orc1) [22] was detectable in the great majority (81%) of both mature and immature gametocytes (Fig 3A). Orc1 is a key component of replication origins in most eukaryotes, and was therefore chosen to represent replication origins in *P. falciparum*. The Orc1 gene in the NF54 strain was epitope-tagged (causing no overt effect on cell viability or gametocyte conversion), and this line was also transfected with a thymidine-kinase-expressing plasmid to ensure that DNA replication could be followed via BrdU incorporation. As expected, Orc1 was mainly (although not exclusively) localised to the nucleus (Fig 3B,C) and there was no clear evidence for the protein changing its distribution or becoming more concentrated in the nucleus as gametocytes matured from stage III to stage V: regardless of stage, it generally appeared either diffused throughout the cell or as concentrated speckles in the nucleus, with a minority of nuclei being fully stained without any speckles (Fig 3A,C). Furthermore, Orc1 was detected identically in the great majority of gametocytes, >80% of which are usually female. Therefore, it was evidently present in both male and female cells, with no distinct subset of cells showing a higher level or a distinct distribution of Orc1.

**Figure 3:**
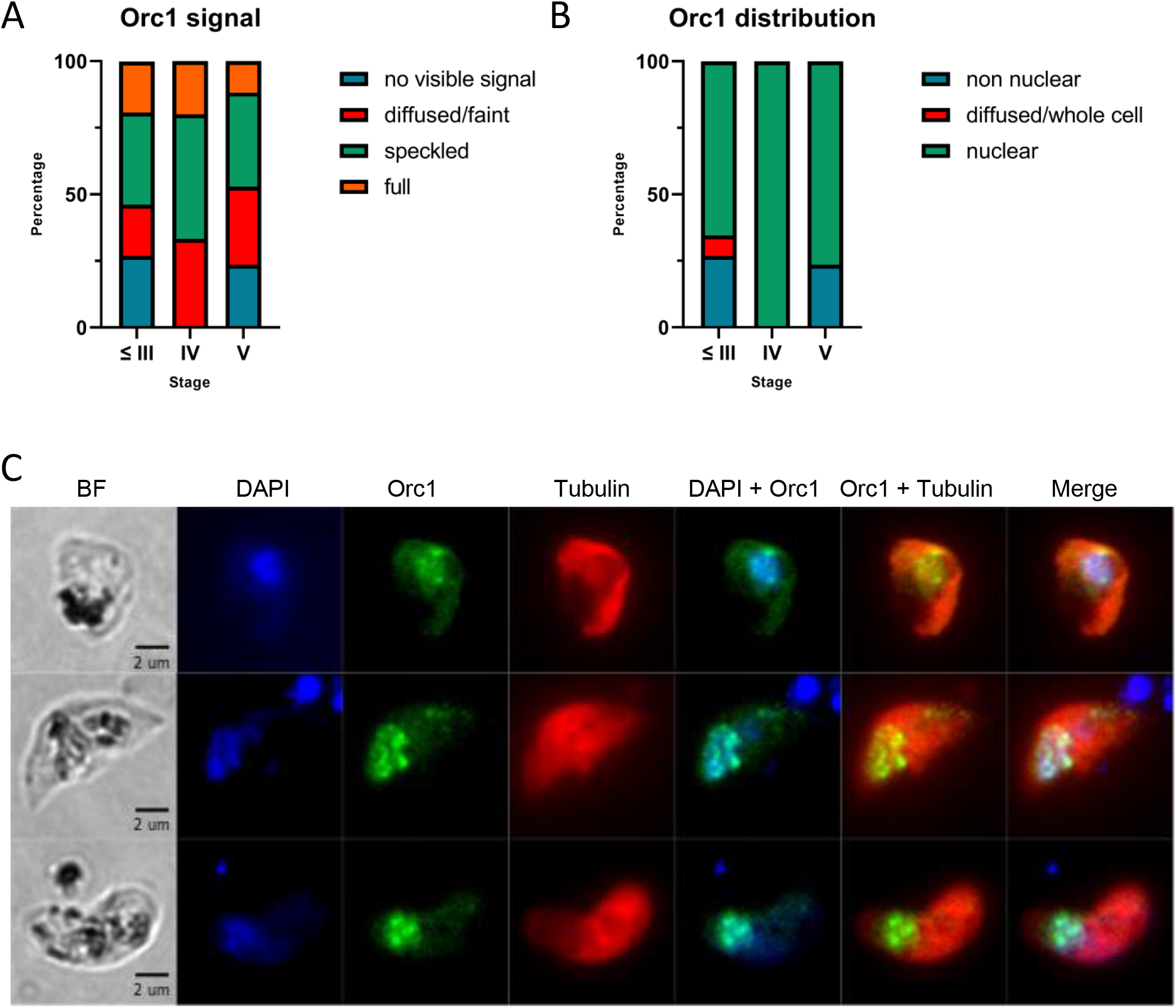
Orc1 measurements in unactivated gametocytes. **A)** Characteristics of Orc1 signal detected in the *P. falciparum* NF54 Orc1-3xHA + pTK-BSD line in gametocytes at different stages (n=15-26 per stage). In total, >80% of gametocytes showed detectable Orc1. **B)** In all cells with detectable Orc1, its distribution was categorised in different gametocyte stages, as in (A). A minority of cells showed Orc1 primarily non-nuclear or distributed throughout the cell, but the majority showed nuclear concentration of Orc1. **C)** Representative images of Orc1 detected in gametocytes at Stage ≤III, IV and V.

### DNA replication begins within 5 minutes of triggering exflagellation

Having established that although replication origin components are present well in advance of triggering, all DNA replication in male gametogenesis must nevertheless occur *after* triggering, we set out to follow this process in detail.

Mature gametocytes were exposed to BrdU, activated via the well-established stimuli of temperature, pH change and xanthurenic acid, then fixed after 0, 5, 10 and 20 minutes to measure DNA synthesis via ELISA and immunofluorescence. At the population level, DNA synthesis clearly commenced within 5 minutes and continued, approximately linearly, through to 20 minutes. During this time, additional α-tubulin also appeared (Fig 4A and Supp. Fig 1). Consistently, at the cellular level, the great majority of male cells that were competent to replicate had started to incorporate BrdU within 5 minutes (Fig 4B), although only a minority of cells completed the process this quickly. Exflagellating cells were scarce at 5 minutes, steadily increasing to the expected proportion of ∼15% by 20 minutes (Fig 4B). Population-level DNA synthesis likewise continued through to 20 minutes (Fig 4A). The majority of the population, representing ∼85% female cells, rounded up upon triggering but did not replicate, as expected, while a small proportion of cells were evidently dead or inviable and remained crescent-shaped after activation (Fig 4B).

**Figure 4:**
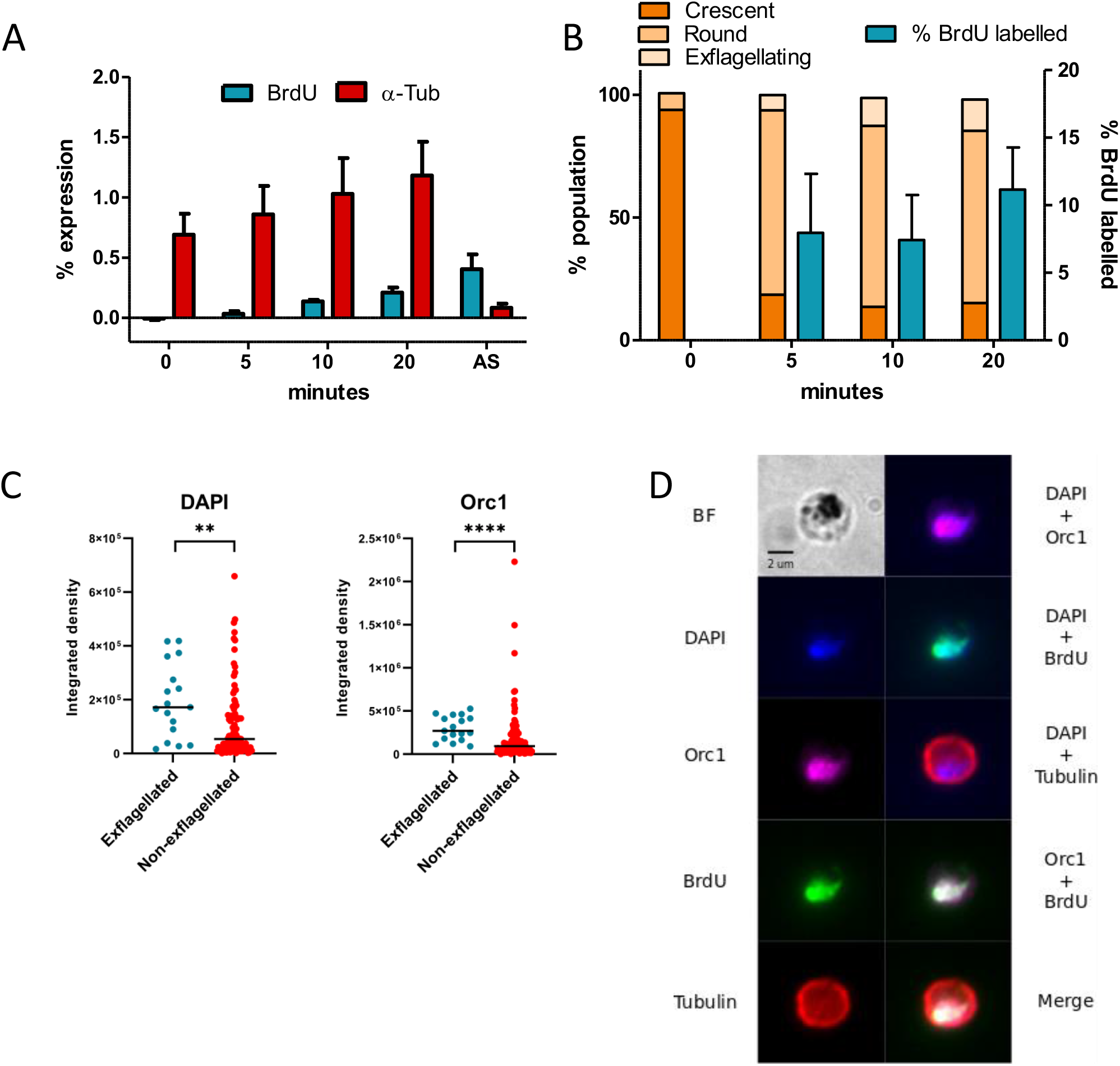
DNA synthesis and Orc1 measurements in activated gametocytes. Following addition of BrdU, exflagellation was triggered in mature gametocytes on day 15 *via* a reduction in temperature and the addition of ookinete media. BrdU incorporation was analysed during the course of exflagellation at 5, 10 and 20 min time points, using the same methods as in **Figure 2**. **A)** Population-level ELISA measurement of DNA synthesis (BrdU) and α-tubulin levels (α-tub). An asexual culture was included as a control. ELISA data represent the mean of at least three independent replicates. **B)** Single-cell-level immunofluorescence images (100 cells per timepoint) were categorised for cellular morphology (left axis) and for BrdU labelling (right axis). Data represent three independent counts of 100 parasites per timepoint. **C)** DAPI and Orc1 signal intensities in exflagellated (BrdU-labelled) versus non-exflagellated gametocytes. Intensities were significantly higher in exflagellated cells (p = 0.0034 for DAPI and p < 0.0001 for Orc1). **D)** Representative image of Orc1 localisation in a gametocyte 30 minutes post exflagellation with BrdU present.

The Orc1 protein was also examined up to 30 minutes after activation, when all viable cells should theoretically have completed the process, Orc1 could be detected in 35% of remaining cells (n=41 of 118 cells), but only 15% (n=17) showed BrdU incorporation. These were classified as male cells that had not yet divided after a full exflagellation event. As expected, the DNA signal as quantified by DAPI staining was much higher in these cells than in those that did not replicate after triggering (i.e. primarily 1n female cells), confirming that they had undergone genome replication, and the Orc1 signal intensity was similarly much higher (Fig 4C). Furthermore, the Orc1 signal was now highly colocalised with the BrdU and DAPI signal, suggesting that Orc1 had become concentrated in the nucleus (or degraded in the cytoplasm) as these male cells underwent replication (Fig 4D).

### Cytokinesis of male gametes can occur without preceding DNA replication

During the timecourses described in Figure 4, we categorised the cellular morphology and nuclear morphology of all male cells – identified by BrdU incorporation – at each timepoint after triggering. Representative images of each morphology are shown in Figure 5A and Supp Figure 1.

**Figure 5:**
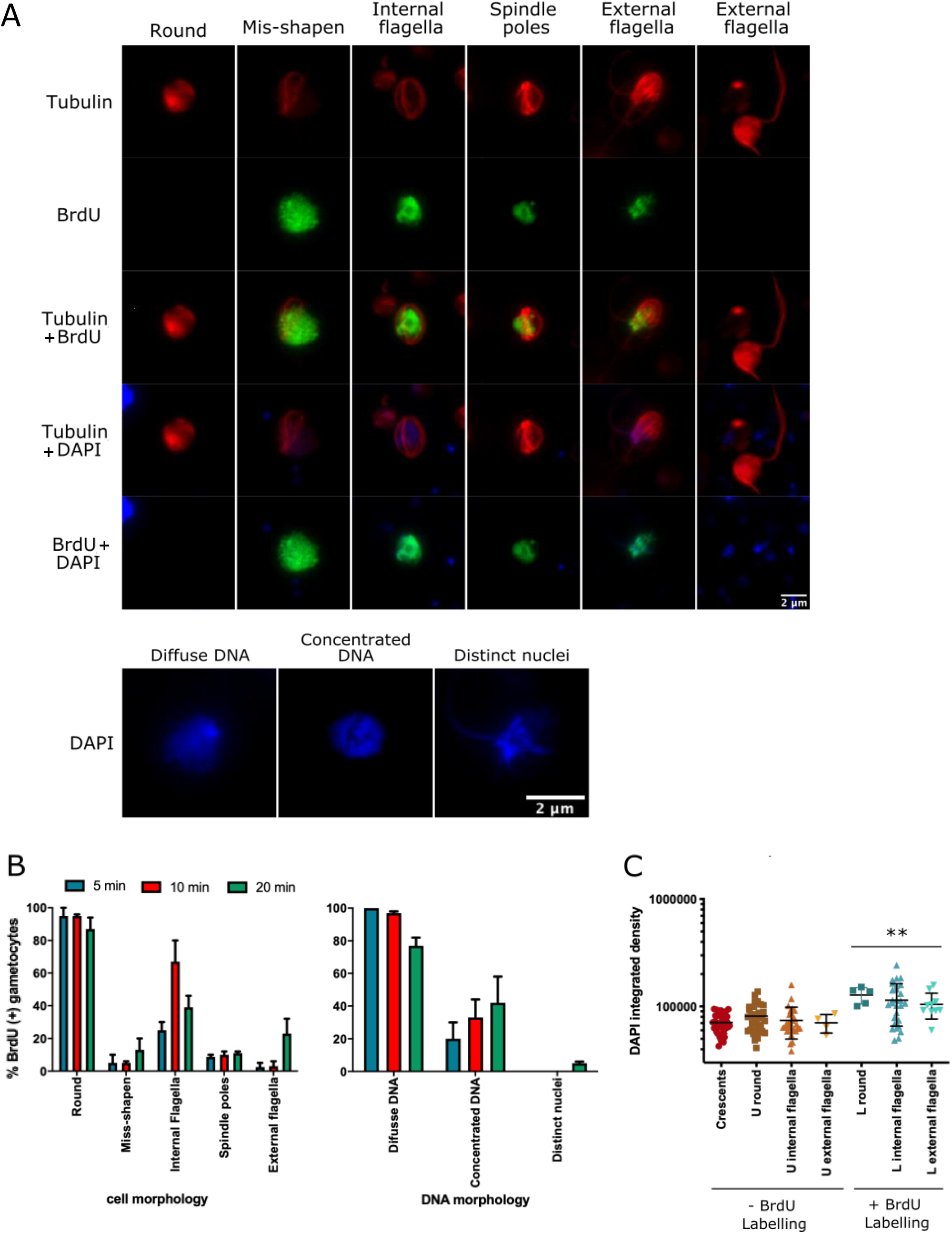
Cellular dynamics of replication and cytokinesis. **A** Representative images of different morphologies of tubulin and DNA seen in male gametocytes. Exflagellating parasites both with and without DNA replication (i.e. BrdU incorporation) are shown. **B)** Parasites incorporating BrdU were analysed for cellular and DNA morphology at 5, 10 and 20 min time points. Data are based on a count of 50 parasites at each time point (note that parasites can appear with more than one feature) and two independent replicates. **C)** The DAPI signal in unactivated parasites, and in activated parasites with and without BrdU incorporation, was quantified using image J. For parasites - BrdU labelling, n = 35 crescents, 37 round, 28 internal flagella, 4 external flagella; for parasites + BrdU labelling, n = 5 round, 28 internal flagella, 10 external flagella. Differences between the amounts of DAPI staining seen in parasites of each morphology +/- BrdU incorporation were assessed by 1-tailed t-test.

Having transformed from their initial crescent-shaped morphology, the great majority of male cells remained round from 5 to 20 minutes, but a growing proportion – reaching ∼10% by 20 minutes – became misshapen with disordered tubulin, possibly indicating aberrant events (Fig 5B). Amongst the round cells, three distinct forms of tubulin were distinguishable: long flagella initially appeared trapped inside the cell (peaking at 10 minutes) and then began to escape outside the cells (peaking at 20 minutes), while ∼10% of cells throughout the timecourse also showed tubulin organised with the appearance of spindle poles, possibly representing hemi- or mitotic spindles, phenomena that are under active investigation in *Plasmodium* schizogony [23]. These suggested that active genome separations were occurring at all time-points.

Alongside these cellular morphologies, nuclear morphology was assessed via DAPI staining (Fig 5A, B). The great majority of cells at all times contained a ball of diffuse DNA, within which more concentrated areas of DNA appeared in ∼50% of cells by 20 minutes. Only a few cells were captured with clearly distinct, condensed nuclei at the 20-minute timepoint, suggesting that this stage is very short-lived and appears only just before the mature gametes separate.

Strikingly, it became apparent that a significant number of cells had clear flagella (diagnostic of male gametogenesis) but no BrdU incorporation (Fig 5A, Supp Fig 1), suggesting that DNA synthesis had not occurred. To confirm this, we quantified the DAPI staining, which scales approximately with DNA content, in cells that were positive or negative for *de novo* DNA synthesis (Fig 5C). Indeed, DNA contents were the same in all cells lacking BrdU incorporation – including untriggered crescent-shaped cells, a large number of rounded cells that must be predominantly females with 1n genomes, and also cells that showed flagella and were therefore presumably male, but lacked BrdU incorporation. By contrast, all cells that incorporated BrdU had higher DNA contents, regardless of whether or not they had yet developed flagella.

### DNA replication dynamics differ in male gametogenesis versus asexual schizogony

To measure the dynamics of DNA replication in male gametogenesis at the single-molecule level, we used the DNA combing technique that was previously used successfully to measure replication fork velocity and replication origin spacing in asexual schizogony [15]. This involves embedding cells in agarose, dissolving their haemozoin and digesting away all cellular proteins, followed by the agarose matrix, thus releasing very long DNA fibres that can be spread onto sialinized glass at a constant stretching factor. Immunofluorescence on these fibres, as on whole cells, can reveal areas of BrdU incorporation and hence *de novo* DNA replication, allowing quantitative measurements of replication fork velocity and replication origin spacing.

Gametocyte cultures were activated in BrdU, then arrested rapidly on ice after time intervals between 5 minutes and 3 seconds, before extracting and combing the DNA fibres. BrdU was then detected in these fibres by immunofluorescence. Figure 6 shows the previously-established pattern seen after a 10-minute pulse label of asexually replicating cells: discrete tracks of newly-replicated DNA, each corresponding to a single replication fork that was active during the pulse label. Remarkably, it was not possible to measure similar tracks from replicating gametocytes, even after a pulse label of just 3 seconds. The great majority of DNA fibres appeared fully replicated for their entire length, usually several hundred kb (Fig 6). Rare fibres were seen with breaks in the BrdU labelling (Fig 6, see ‘distinct tracks’), but these were too scarce to measure meaningful inter-origin distances or average fork velocities. We concluded that it was not technically possible to trigger and then arrest DNA replication in gametocytes before the great majority of DNA fibres had been fully replicated over distances of several hundred kb. This also confirmed that the labelled DNA fibres were predominantly from male gametocytes and not from any contaminating asexual cells (such cells were extremely rare after 15 days of gametocyte culture, and they would appear with a very different pattern of DNA fibre labelling, dominated by discrete tracks).

**Figure 6:**
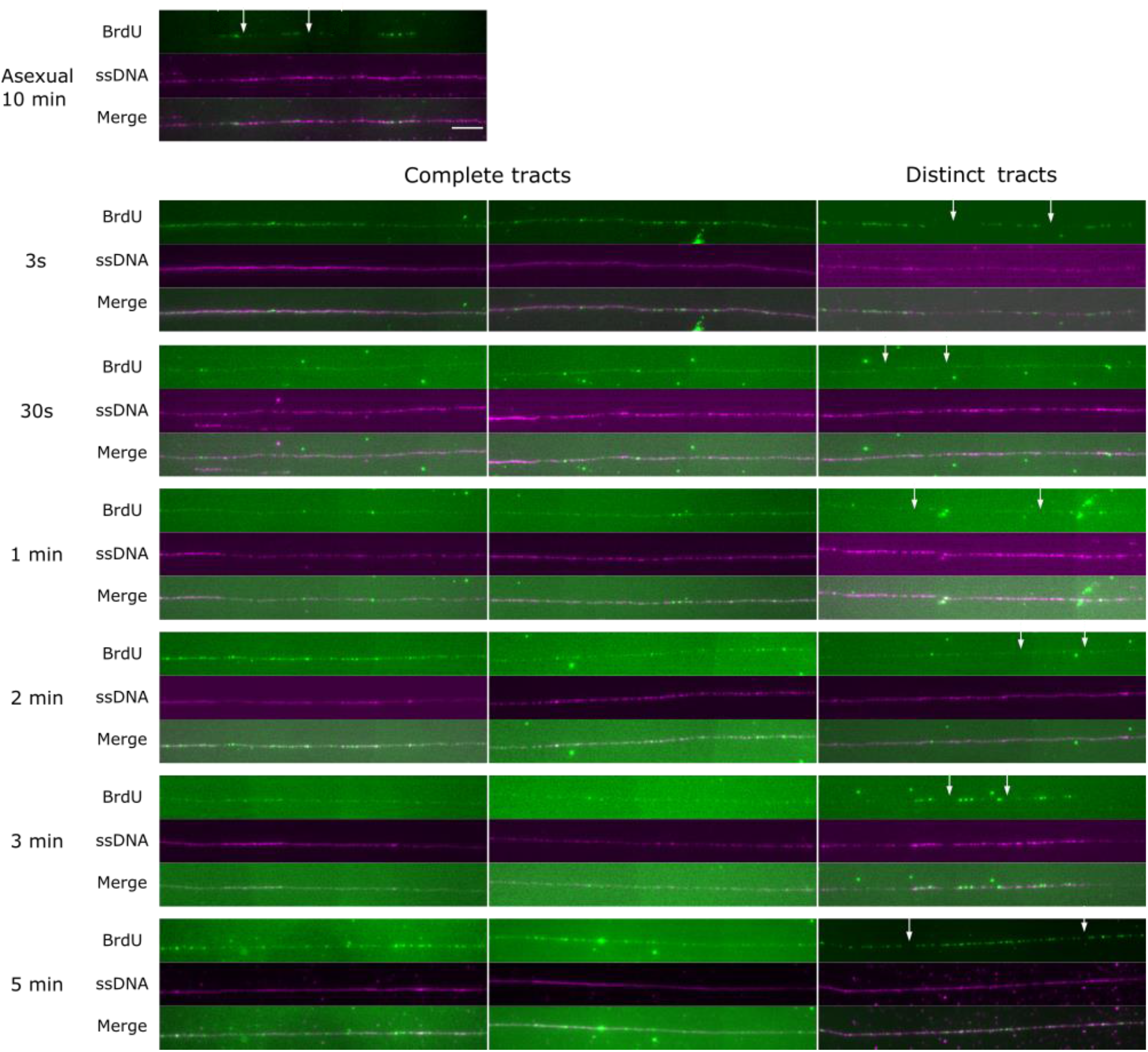
Single-molecule DNA replication dynamics during male gametogenesis. Single-molecule DNA fibres labelled with BrdU for between 3s and 5mins after triggering exflagellation in gametocytes, compared to DNA fibres from asexual erythrocytic parasites after 10 mins of BrdU incorporation. Representative DNA fibres with distinct and continuous BrdU-labelled tracks are shown. Scale bar is 5µm (10 kb). White arrows highlight some of the breaks in BrdU labelling that are seen between distinct tracks.

We next attempted to measure inter-origin spacing with two consecutive pulses of different modified nucleosides. If cells are allowed to switch between two different pulse labels in the course of replication, then two-coloured DNA fibres are detected, hence the distance between adjacent origins can be measured even if several adjacent replication bubbles have terminated before the end of the labelling period [15]. Gametocytes were exposed to EdU in an attempt to load the cells with a different nucleotide for initial replication, then washed, switched into BrdU, triggered and quickly arrested. The result was identical to that shown in Figure 6: all fibres still appeared fully replicated in BrdU. Therefore, gametocytes prior to triggering added no detectable amount of exogenous nucleosides to their nucleotide pools. This result is consistent with prior work that attempted to pre-load gametocytes with radiolabelled hypoxanthine for 90 minutes before triggering, and was likewise unable to detect any of the label after triggering [24].

## DISCUSSION

This study used state-of-the-art molecular methods to examine a highly unusual cell-biological process: genome replication in *P. falciparum* male gametocytes. Our findings illuminate this biologically-unique process at a new level of spatial and temporal resolution.

We first established that there is no detectable DNA replication at any point in the 14-day development of male gametocytes prior to gametogenesis. It was previously suggested in early studies – from relatively low-resolution fluorometry on gametocyte nuclei – that some replication occurred in the arrested gametocyte, raising the DNA content to ∼1.5n [4, 5, 8]. This could putatively explain the rapidity of the first replicative cycle in gametogenesis. However, *de novo* DNA replication was never observed in maturing gametocytes, meaning that all replication must occur during the brief window of gametogenesis. Furthermore, we found no evidence that nucleotide pools can be substantially pre-loaded, so 8 genomes-worth of nucleotides must be synthesised as well as assembled into DNA during the time-window of gametogenesis. We also examined the Orc1 protein that is a component of replication origins, and found that it was already present in early gametocytes and was not detectably upregulated or relocalised as gametocytes matured. Surprisingly, it also appeared to be equally abundant in female and male cells, since there was no distinct population high-Orc1-expressing cells, as would be expected if the minority male population contained a high level of this replication factor in readiness for gametogenesis. After activation, however, we did observe elevated levels of highly nuclear-concentrated Orc1 in active male cells. This suggested that replication proteins, as well as nucleotides, may be produced and re-localised during the active process of gametogenesis itself.

Secondly, after triggering gametogenesis we monitored DNA replication at three levels: the population, single cell and single molecule. At the population level, we observed that newly replicated DNA appears within 5 minutes of triggering, consistent with previous reports [4, 5, 8], and continues to accumulate through to 20 minutes. At the cellular level, the great majority of viable male cells were synthesising DNA within 5 minutes, suggesting that triggering was quite synchronous. Finally, at the single-molecule level, it became clear that DNA replication initiates extremely rapidly in at least some male gametocytes, since large numbers of newly replicated DNA fibres could be detected even if the process was halted within a few seconds of triggering. By contrast, the time taken to complete gametogenesis was very variable: exflagellating cells initially appeared within 5 minutes, and continued to appear in greater numbers through to 20 minutes, indicating that gametogenesis could take anything from 5 to >20 minutes in total.

By observing the morphology of replicating DNA in large numbers of cells across the course of gametogenesis, we confirmed that there is no apparent karyokinesis between rounds of DNA replication – in sharp contrast to the steadily increasing numbers of discrete, condensed nuclei that appear during erythrocytic schizogony [13]. Discrete nuclei were only ever seen at the final stages of exflagellation and it was very rare to capture such cells, emphasising the extreme brevity of this stage. Nevertheless, there was some evidence for the separation of genomes after each replication. Areas of denser DNA within the mass of diffuse DNA became visible over time, perhaps representing partially-condensed nuclear masses, and ∼10% of cells throughout gametogenesis also contained tubulin in the form of spindles, consistent with regular rounds of chromosomal separation. This observation is consistent with other ultrastructural studies of *P. yoelii* [25] and *P. falciparum* [26] gametogenesis, and with two very recently-published studies using either live-imaging or expansion microscopy [27, 28].

Strikingly, there appears to be no DNA replication checkpoint prior to cytokinesis in gametogenesis. Cells were regularly observed exflagellating without any evidence of *de novo* DNA replication, and indeed these cells had total DNA contents comparable to the 1n content of untriggered cells. Thus it is evidently possible for the cytokinesis programme to proceed despite a complete failure to replicate the genome. Why DNA replication should frequently fail remains unknown. Because exflagellating cells with condensed nuclei were captured so rarely, it was impossible to assess what happens to a single unreplicated genome: is it partitioned into a single gamete, leaving others as anucleate zoids, or is it separated aberrantly into eight fragmentsã Furthermore, it seems likely that exflagellation could still occur if replication produced fewer – or more – than 8 full genomes: although this appears to be the norm in all *Plasmodium* species, there are anecdotal reports in the early literature of partial replication events yielding some nucleated and some zoid gametes [18] (and the number of microgametes produced by the equivalent process in the related species *Toxoplasma* is very variable [29]).

The observation that ∼10% of cells were ‘misshapen’ by 20 minutes post-triggering, containing disordered tubulin and probably representing failed cells, certainly suggests a high failure rate, at least in our experiments, and this is also consistent with observations in the early literature. Importantly, our experimental system may not be as stringent and well-controlled as the *in vivo* situation: inside a mosquito, the synchrony, fidelity and success rate of gametogenesis might all be greater, and *P. berghei* gametocytes that have developed in a mouse rather than a petri plate tend to behave more synchronously [24]. All single-molecule measurements, meanwhile, were limited by the accuracy with which parasites could be triggered *in vitro* and then chilled within a few seconds. Nevertheless, the broad conclusions from these *in vitro* experiments are likely to apply *in vivo*, since aberrant events, as well as zoid gametes and cells producing fewer than 8 gametes, have all been observed in *P. berghei* gametogenesis [5, 18].

At the single molecule level, we were unable to measure replication parameters accurately because the resolution of DNA combing proved limiting – although the technique was quite adequate for measuring replication dynamics in schizogony. In gametocytes, after a mere 3-second label the majority of DNA fibres appeared completely replicated. Completing a round of replication within 3 seconds using parameters measured in schizogony (origin spacing 65kb, replication speed 1.2kb/min), would necessitate replication speeds of ∼600kb/minute, or origin spacings of ∼120bp, which are not realistic. However, replication is an enzymatic process that probably cannot be halted instantly on ice, so some run-on replication beyond 3 seconds probably occurs. Nevertheless, each round of replication could plausibly take as little as a minute, since some cells can reach exflagellation successfully within 5 minutes, and mitotic spindles have been reported to appear in *P. yoelii* gametocytes within a minute of triggering [25]. The resultant parameters – fork speeds of 32kb/minute or origins every 2.4kb – would still be unprecedented, but such a density of origins is at least physically feasible. (It is perhaps surprising that there was no clear increase in Orc1 protein as gametocytes matured, despite the fact that male gametocytes must presumably lay down a very high density of origins on their genomes prior to triggering replication.) Alternative explanations for extremely fast replication, such as re-replication, cannot be strictly excluded but we found no evidence for re-replication in the form of variable-density BrdU labelling of DNA fibres.

The extreme speed of replication in male gametogenesis is probably driven by evolutionary pressure to complete a sexual cycle before parasites are digested along with the blood meal or attacked by human immune factors within it. Our data support the idea that this has enforced a tradeoff of speed versus fidelity, leaving *Plasmodium* gametocytes particularly vulnerable to replication failures, particularly since cell cycle checkpoints seem to be absent. Compounds that inhibit replication or damage DNA could therefore block male gamete production in mosquitoes – and indeed, it may be possible to poison the replicative capacity of gametocytes while still in the human bloodstream, thus blocking subsequent transmission [30]. Many anti-cancer therapies are informed by an analogous approach in cancer cells, exploiting vulnerabilities in their rapid cell division rate. The parameters described in this study, and the methods for measuring DNA replication in gametocytes, may be very useful in future studies of transmission-blocking drugs.

## METHODS

### Parasite culture and transfection

The NF54 strain of *P. falciparum* (gift from Prof Baum, Imperial College London; also available from MR4 repository, www.beiresources.org) was used for all experiments. Parasites were maintained *in vitro* in human O+ erythrocytes at 4% haematocrit in RPMI 1640 medium supplemented with 25mM HEPES (Sigma-Aldrich), 0.25% sodium bicarbonate, 50 mg/L hypoxanthine (Sigma-Aldrich), 0.25% Albumax (Invitrogen) and 5% heat-inactivated pooled human serum, using standard procedures [31]. Transfection with a plasmid carrying the HSV thymidine kinase gene was conducted as previously described [21]. The Orc1 gene (PF3D7_1203000) in *P. falciparum* NF54 was C-terminally tagged with 3xHA by selection-linked integration (SLI) [32]. Briefly, ring stage parasites were transfected with 100 µg of pSLI plasmid (with a human DHFR selectable marker cassette) using a Biorad Gene Pulser XL electroporation system. For homologous recombination, the 3’ 505 bp of the Orc1 gene (excluding the stop codon) was cloned upstream of 3xHA, a skip peptide, and a neomycin positive selection marker. Transfectants were initially selected with 5 nM WR99210 followed by 400 µg/mL of G418, and were confirmed by PCR and western blot. To allow the transfectants to utilise thymidine analogues (e.g., bromo-deoxyuridine, BrdU), ring stage transfectants (*i*.*e*., *P. falciparum* NF54 Orc1-3xHA) were transfected with 100 µg of blasticidin (BSD)-selectable plasmid carrying the HSV thymidine kinase gene with a *hsp86* promoter and a *P*.*berghei dhfr* 3⍰. terminator. Transfectants were selected with 2.5 µg/mL BSD. Successful transfectants (*i*.*e*., *P. falciparum* NF54 Orc1-3xHA + pTK-BSD) were confirmed by plasmid rescue and immunofluorescence after BrdU incorporation. Final transfectants were maintained in 5 nM WR99210 and 2.5 µg/mL BSD.

### Production of gametocytes

Gametocyte production methods were consistent with those described by Delves *et al*. [33]. Briefly, asexual feeder cultures were maintained at 4% haematocrit < 5% parasitaemia in serum-only culture media. For gametocyte culture setup, predominantly ring-stage cultures were seeded at 2% rings (<0.5% other stages), 4% haematocrit with complete media containing 10% serum and 0.25% albumax. Spent media was removed and replaced with pre-warmed (37 °C), fresh media, daily for 14 days. Gametocyte cultures were placed on a heat block, set at 38 °C, to limit temperature variations during the media exchange. Only serum that had been batch tested for viable gametocyte production was used. All NF54-pTK asexual and gametocyte cultures were maintained with 5nM WR. On days 5-7 N-Acetyl-D-glucosamine (20mM) was added (i.e. 48 h treatment) to remove asexuals. Cultures with >0.2% exflagellation were used for experiments.

### Gametocyte purification and exflagellation for ELISA and IFA assays

Gametocytes were purified using a NycoPrep density gradient. Gametocyte culture volume was reduced to 10 ml, layered onto 10ml nycoprep (Progen), and centrifuged at 800g for 20 mins. The gametocyte interface was removed, washed twice with RPMI and resuspended in serum-only complete medium. The centrifuge and all solutions and were pre-warmed to 37oC to prevent premature exflagellation.

Following gametocyte purification, gamete formation was triggered *via* a drop in temperature and the addition of 50µl ookinete media containing 100 µM xanthurenic acid per 800 µl of parasite suspension. (Ookinete media composition: 8.15 g RPMI with L-glutamine, phenol red, HEPES, without sodium bicarbonate; 1g NaHCO3; 25 mg hypoxanthine, adjusted to pH 7.4 using NaOH). Cultures were used only if they displayed at least 0.2% exflagellation. At 5, 10 and 20 minute intervals samples were taken in parallel for IFA, ELISA and SYBR green-based measurement of parasitemia (Malaria SYBR green fluorescence assay (MSF)). For IFA, samples were immediately fixed in 4% formalin.

### BrdU labelling for ELISA and IFA assays

DNA replication was analysed at both the population level (ELISA) and the single-cell level (using immunofluorescence assays (IFAs)) in parallel. To measure nascent DNA synthesis, the incorporation of 5-bromo-2’-deoxyuridine (BrdU) was evaluated during gametocytogenesis (‘pre-triggering’) at days 8, 10, 12 and 14. Cultures were incubated with 100µM BrdU for 6 h prior to purification. For the evaluation of DNA synthesis during gametogenesis (‘post-triggering’) the BrdU label (100µM) was added after purification, immediately prior to exflagellation induction.

### Gametocyte-to-gamete ELISA and MSF assay

Aliquots of parasites taken at each time point were distributed (in triplicate) into two replicate 96 well plates. At least 2.5×10^5^ exflagellation centres were plated per well. One plate was used immediately for estimation of parasitaemia via genome staining with SYBR green, as described previously [21]. In brief, SYBR Green I/ MSF lysis buffer was added, incubated for 1 h at RT and analysed using the GloMax multidetection system (Promega). Parasitaemia was used to normalise parasite density between samples and replicates. The second plate was airdried at 37 °C overnight for the ELISA. Once dried, the ELISA plate was fixed with 4% formaldehyde/PBS for 2 min, followed by 50% acetone/50% methanol for 2 min, and then blocked for 1 h with 1% BSA/PBS. Primary antibodies were then added, mouse anti-BrdU/nuclease (1:100) (Amesham) or mouse anti α-tubulin (Sigma)/1% BSA/PBS (1:500) and incubated for 2 h at 37 °C. Tubulin was used as a control to allow visualization of flagella formation and the progression through gametogenesis in relation to DNA synthesis. Following 3 washes with PBS, the secondary antibody, anti-mouse HRP (1:5000) was added and incubated for 1 h at RT. After a further 3 washes with PBS, samples were developed with 50µl/well of Single-Component TMB Peroxidase EIA Substrate Kit (BioRad) for 5-10 min. The reaction was terminated by the addition of 0.6M sulphuric acid. The plate was analysed using the GloMax multidetection system (Promega).

ELISA readings were normalised to the starting parasite density, as described above, and also blank-subtracted using readings from a parallel culture that was not exposed to any BrdU. In addition to providing a background reading as a blank, this parallel experiment confirmed that gametogenesis was not impeded by the addition of BrdU (i.e. tubulin production and exflagellation appeared to progress normally).

### Gametocyte-to-gamete immunofluorescence assay

In parallel with the 96 well plates set up for ELISA and SYBR green-based parasitemia estimates, as described above, aliquots from each time point were transferred to 12 well, poly-L-lysine coated slides for immunofluorescence assays. Following overnight storage (or up to 3 days) in a humidified chamber at 4°C the slide was blocked for 30 min in 1% BSA/1xPBS, permeabilised with 0.1% Triton-X100/1x PBS for 15 min and washed 3 times in 1x PBS. Primary antibodies were then added: mouse anti-BrdU/nuclease (1:100) (Amersham) and rabbit anti-αTubulin/1% BSA/1x PBS (1:500) (Abcam Ab52866) and incubated for 1 h at RT. α-tubulin was used as a control to allow visualization of flagella formation and the progression through gametogenesis in relation to DNA synthesis. Following 3 wash steps in 1x PBS, secondary antibodies were added and incubated for 45 min at RT: anti-mouse Alexa488-conjugated (1:500) and anti-rabbit Cy3-conjugated (1:500) (Molecular Probes). After washing (3x in 1x PBS) coverslips were mounted using Vectashield+ DAPI (Molecular Probes) and left to set overnight at 4°C. DAPI was included to confirm DNA location and morphology. Slides were viewed and images were collected using an EVOS fluorescence microscope (Thermofisher scientific).

### Orc1 immunofluorescence assay

A total of 20 mL of NF54-Orc1-3xHA + pTK-BSD gametocyte culture was prepared, harvesting 10 mL at day 9 and 10 mL at day 14. Gametocytes were purified using prewarmed 55% Nycodenz (Alere Technologies AS) in RPMI from 100% stock (Nycodenz 27.6% (w/v), Tris HCl 5 mM, KCl 3 mM, EDTA 0.3 mM). Purified gametocytes were fixed in 4% formaldehyde and mounted on poly-L-lysine coated slides. Fixed cells were allowed to settle on the slide at least overnight in a humidified chamber at 4°C. Slides were washed 3 times in 1xPBS and were blocked for 1 hour in 1% BSA/1xPBS with 0.1% Tween-20. Slides were incubated at room temperature in the following primary antibodies at 1:500 dilution in 1% BSA/1xPBS: rat anti-HA (Roche) and rabbit anti-αTubulin (Abcam). Slides were then washed 3 times in 1% BSA/1xPBS and were incubated in 1:1000 dilution (in 1% BSA/1xPBS) of the following secondary antibodies: Alexa488-conjugated goat anti-rat (Molecular Probes) and Alexa647-conjugated goat anti-rabbit (Molecular Probes) for 1 hour at room temperature. Secondary antibodies were washed off 2x with 1% BSA/1xPBS, and were incubated in 2 ug/mL of DAPI for 5 minutes. Coverslips were mounted using Prolong Diamond Antifade (Molecular Probes) and were allowed to set overnight at room temperature prior to imaging. Gametocytes were staged as ≤ III, IV, and V by morphology. Orc1 and DAPI signal intensities for individual gametocyte were measured using ImageJ. Statistical analyses (Kruskall-Wallis and Mann-Whitney U tests) to compare signal intensities in different gametocyte stages were done using GraphPad PRISM.

For gametocyte exflagellation in BrdU, gametocytes were purified from 20 mL of gametocyte culture and were resuspended in warm gametocyte media supplemented with 100 µM of BrdU. Ookinete media was added to the gametocytes, which were fixed in 4% formaldehyde 30 minutes post-exflagellation. Immunofluorescence staining was done as above with the addition of mouse anti-BrdU (Merck) primary antibody at 1:500 in 1 U/mL DNAse I (Amersham) in water and Alexa594-conjugated goat anti-mouse secondary antibody (Molecular Probes).

### Preparation of agarose plugs for combing of *P. falciparum gametocyte* DNA

For labelling of gametocyte cultures, 100 µM 5-bromo-2’-deoxyuridine (BrdU) was added to the culture before triggering of exflagellation. Exflagellation was triggered to begin DNA replication by the addition of ookinete media (1:1 ratio) at room temperature of 21°C at timepoint 0. Replication was stopped at designated timepoints (0, 0.5, 1, 5 mins, etc.) by adding ice cold PBS and keeping on ice at all times. Culture was washed in ice cold PBS and resuspended in 0.1% saponin for 5 mins. The culture (1×108 gametocytes/ 100 µl PBS) was then washed three times in ice cold PBS at 4°C and resuspended in 100µl PBS containing 1% low melting point agarose (Dutscher) to make plugs as previously described [15]. Each plug was incubated in 1 ml Proteinase K buffer (10 mM Tris HCl pH 7.5, 50 mM EDTA, 1 % N-lauryl-sarcosyl, 2mg /ml proteinase K in dH20 at 45°C for two days with fresh Proteinase K buffer added on the second day. The complete removal of any digested proteins or degradation products was effected by five, 1h washes with TE50 buffer (50 mM EDTA, 10 mM Tris HCl pH 7.5) with slight agitation of plugs. Plugs were then stored in TE50 buffer at 4°C or used immediately for combing.

### DNA Molecular Combing

DNA combing was performed as previously described [15] using silanised CombiCoverslips (Genomic Vision) and the Molecular Combing System (Genomic Vision), producing a constant stretching factor of DNA fibres of 2kb/ µm.

### Detection of BrdU in DNA fibres

Combed DNA on coverslips was baked overnight at 65°C, denatured with 1M NaOH for 20’ and washed three times in PBS. Coverslips were washed four times in PBS and blocked with 1% BSA and 0.1% Tween 20 in PBS. Immuno-detection was done with antibodies diluted in PBS, 1% BSA, 0.1% Tween 20, at room temperature for 1h, covered with a second coverslip to prevent dehydration. Each step of incubation with antibodies was followed with three washes in blocking buffer. Primary immuno-detection was done with rat anti BrdU BU1/75 (ICR1) antibody (1:100 dilution, Abcam) and mouse anti ssDNA (clone16-19) antibody (Millipore, 1:300 dilution). The secondary antibodies (Molecular Probes) were goat anti-rat coupled to Alexa 488 (1:500 dilution) and goat anti-mouse coupled to Alexa 594 (1:500 dilution). Coverslips were washed three times in PBS and mounted using 20µl Prolong Diamond Antifade (Molecular Probes), set overnight at room temperature.

### Image Acquisition and Processing

Image acquisition was via a Nikon Microphot SA microscope equipped with a Qimaging Retiga R6 camera. Images were acquired with a 100X oil objective where 1 pixel corresponds to 64.80 bp (DNA stretching factor 2kb/µm). Observation of long DNA fibres required the capture and assembly of adjacent fields. Replication tracts and fibre lengths were measured manually using ImageJ software.

---

**SUPP FIGURE 1.**
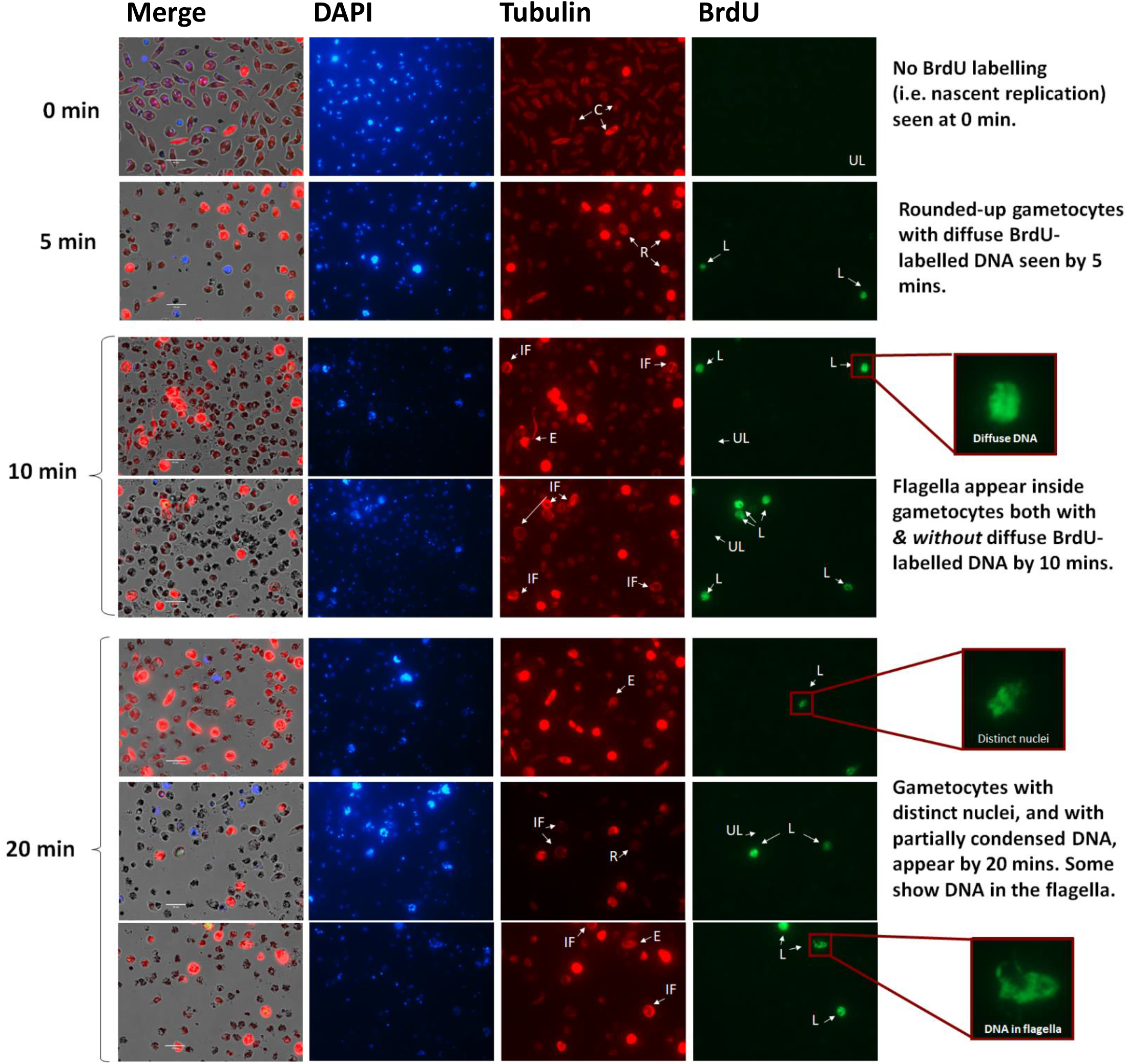
Parasite morphology during 20 minute timecourse after triggering of gametogenesis. Crescent shaped (C), Round (R), Round with internal flagella (IF), Exflagellating (E), BrdU-labelled (L), no BrdU incorporation i.e. unlabelled (UL). Males make up 13-17% of the population; the remaining majority of females also round up but never show BrdU labelling.

